# Selection for synchronized replication of genes encoding the same protein complex during tumorigenesis

**DOI:** 10.1101/496059

**Authors:** Ying Chen, Ke Li, Xiao Chu, Lucas B. Carey, Wenfeng Qian

## Abstract

DNA replication alters the dosage balance among genes; at the mid-S phase, early-replicating genes have doubled their copies while late-replicating genes have not. Dosage imbalance among proteins, especially within members of a protein complex, is toxic to cells. Here, we propose the synchronized replication hypothesis: genes sensitive to stoichiometric relationships will be replicated simultaneously to maintain stoichiometry. In support of this hypothesis, we observe that genes encoding the same protein complex have similar replication timing, but surprisingly, only in fast-proliferating cells such as embryonic stem cells and cancer cells. The synchronized replication observed in cancer cells, but not in slow-proliferating differentiated cells, is due to convergent evolution during tumorigenesis that restores synchronized replication timing within protein complexes. Collectively, our study reveals that the selection for dosage balance during S phase plays an important role in the optimization of the replication-timing program; that this selection is relaxed during differentiation as the cell cycle is elongated, and restored as the cell cycle shortens during tumorigenesis.

## INTRODUCTION

The balance hypothesis asserts that the stoichiometric relationship among subunits of a protein complex is essential for the survival and proliferation of cells; the disruption of this relationship perturbs functions of protein complexes and sometimes even causes cytotoxicity (1–8). The balance hypothesis provides a unique framework for understanding a variety of biological phenomena, especially the proliferation rate of aneuploid cells and the fate of duplicated genes. Aneuploidy, defined as a karyotype that is not a multiple of the haploid complement, generates dosage imbalance among genes on different chromosomes. Consistent with the balance hypothesis, aneuploidy often results in a more severe growth defect than a whole genome duplication that keeps the dosage balance among genes (5, 9). Furthermore, the addition of a larger chromosome, which leads to a dosage imbalance among more genes, often results in a greater reduction in fitness (6, 10–13). Gene duplication confers the second type of dosage imbalance, between duplicate genes and singletons. Consistent with the balance hypothesis, genes often reduce their expression soon after duplication (14), through which the dosage balance is restored. Furthermore, genes encoding protein complexes exhibit a higher retention rate after the whole genome duplication so that the dosage balance among subunits is maintained (2, 15).

A probably more prevalent but less studied source of dosage imbalance is caused by DNA replication that occurs each cell cycle. During the DNA synthesis phase (S phase) of a cell cycle, the genome is replicated in a defined temporal order known as the replication-timing program (16, 17). In the middle of S phase, early-replicating genes have doubled their copy number, but late-replicating genes have not, leading to a dosage imbalance between early and late-replicating genes (**Fig. 1A**). Such dosage imbalance likely causes a growth defect especially among genes sensitive to dosage relationship such as those encoding the same protein complex (2, 6). Although acetylated histones (H3K56ac) can incorporate into newly replicated DNA regions and partly suppress the expression of newly replicated genes in yeast (18, 19), this compensatory mechanism cannot completely restore the dosage balance; the mRNA levels of early-replicating genes still exhibited a ~20% increase compared to late-replicating genes during the mid-S phase (18). Consistent with this, a GFP reporter inserted into early-replicating regions in yeast exhibits higher expression (20).

**Figure 1.**
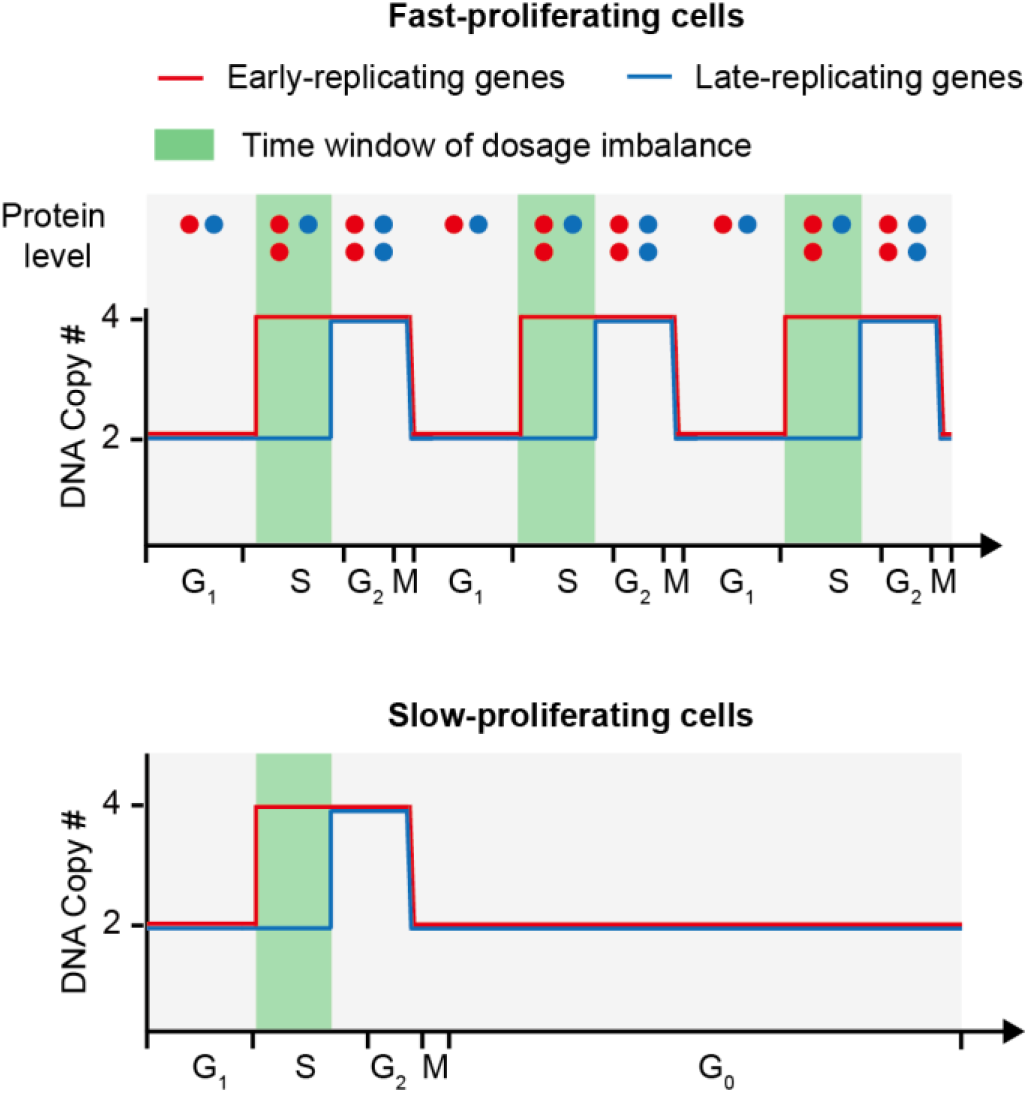
Dosage imbalance between early and late-replicating genes during S phase in fast-proliferating (A) and slow-proliferating (B) cells. The demand for dosage balance during S phase is higher in fast-proliferating cells.

The dosage imbalance during S phase could be severer in mammalian cells where DNA replication lasts longer (~8 hours) each cell cycle. An exacerbating factor is that H3K56ac may not mark newly replicated DNA in mammalian cells (21). Consistently, in mouse embryonic stem cells (ESCs), the transcription rates of *Oct4* and *Nanog* increased by 28% and 50%, respectively, upon DNA replication (22). These data show that replication can cause dosage imbalance during S phase, and suggest that additional mechanisms should exist in mammalian cells to solve the problem. Here, we proposed a hypothesis that the replication of genes encoding the same protein complex is synchronized during S phase so that the dosage balance is warranted. Indeed, we observed a synchronized replication within protein complexes, but surprisingly, only in fast-proliferating cells such as various tumor cells, indicating a convergent evolution towards synchronized replication during tumorigenesis.

## RESULTS

### Genes encoding subunits of the same protein complex tend to replicate simultaneously in HeLa cells

The synchronized replication hypothesis predicts a reduced variation in replication timing among genes encoding the same protein complex. Indeed, genes encoding some protein complexes are replicated almost simultaneously in HeLa cells as exemplified in **Fig. 2A**. To test this prediction at the genomic scale, we retrieved the components of 1,521 protein complexes from the Human Protein Reference Database (23) and the replication-timing program of HeLa cells (24). For each protein complex, we calculated the standard deviation of replication timing of all genes encoding the protein complex (**Fig. 2B**). As a control, we randomly sampled genes from the genome to constitute “pseudo” protein complexes, keeping the number of complexes and the number of subunits in each complex unchanged (**Fig. 2B**). We performed the random sampling 1,000 times. The median of observed standard deviations is significantly smaller than the random expectation (*P* < 0.001, permutation test, **Fig. 2B**), indicating synchronized replication within protein complexes. The same conclusion can be reached when we shuffled among genes encoding protein complexes to constitute “pseudo” protein complexes (*P* < 0.001, permutation test, **Fig. 2C**).

**Figure 2.**
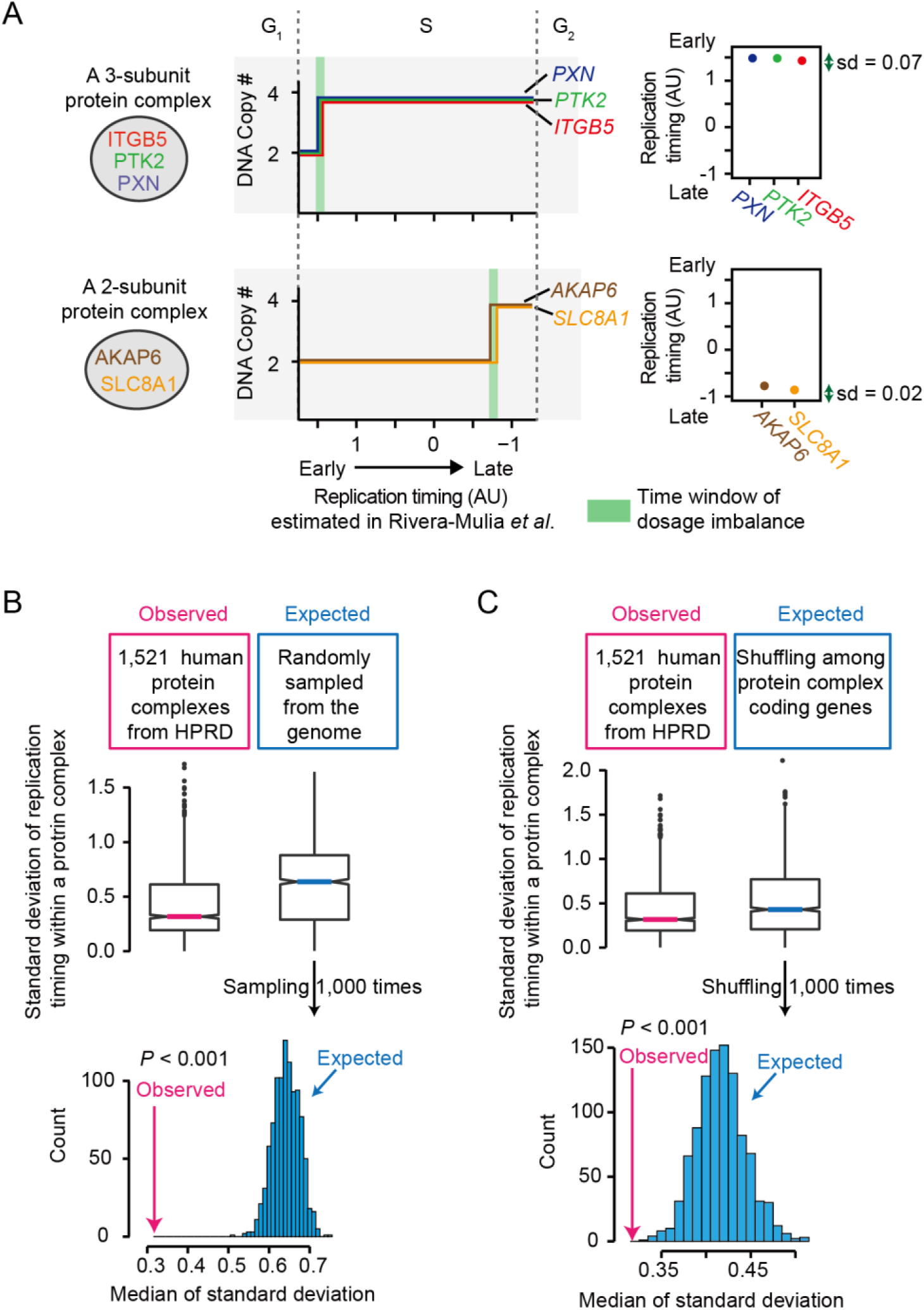
Genes encoding the same protein complex are replicated simultaneously in HeLa cells. (A) Two examples showcase the synchronized replication of genes encoding the same protein complex. The standard deviation of replication timing within a protein complex is shown on the right. (B-C) The observed standard deviation of replication timing within a protein complex is significantly smaller than the random expectation where the protein complex-coding genes were randomly sampled from the genome (B) or shuffled (C). The protein complex information was retrieved from the Human Protein Reference Database (HPRD).

It is worth noting that genes encoding the same protein complex tend to form clusters on chromosomes (25, 26), which are likely to simultaneously replicate because they have similar physical distances to the closest replication origin. To determine if the smaller variation in replication timing within a protein complex is fully explained by such gene clusters, we discarded protein complexes of which at least two subunits are encoded by the genes on the same chromosome. The smaller standard deviation of replication timing within a protein complex remained observed (*P* = 0.002, permutation test, **Fig. S1**).

### Synchronized replication occurs only in fast-proliferating cells

The replication-timing program varies among cell types. To determine if synchronized replication occurs uniformly among various human cells, we retrieved the replication-timing programs previously reported in 17 cell lines/types (24, 27). They include 6 human ESC lines, 5 cancer cell lines, and 6 differentiated cell types such as liver and pancreas cells derived from human ESCs (**Fig. 3A**). The proliferation of ESCs and cancer cells is fast whereas that of differentiated cells is slow (28). These cell lines/types exhibit various levels of synchronized replication within protein complexes (as exemplified in **Fig. 3B**). To assess synchronized replication at the genomic scale, we randomly shuffled genes encoding protein complexes (**Fig. S2**, with three examples shown in **Fig. 3C**). We used the *P* value of the permutation test to infer the level of synchronized replication in each cell line/type and labeled synchronized replication for those with *P* < 0.05 (**Fig. 3E**). Surprisingly, synchronized replication was exclusively observed in the 11 fast-proliferating cell lines (**Fig. 3D-E**, *P* = 8×10^−5^, the Fisher’s exact test).

**Figure 3.**
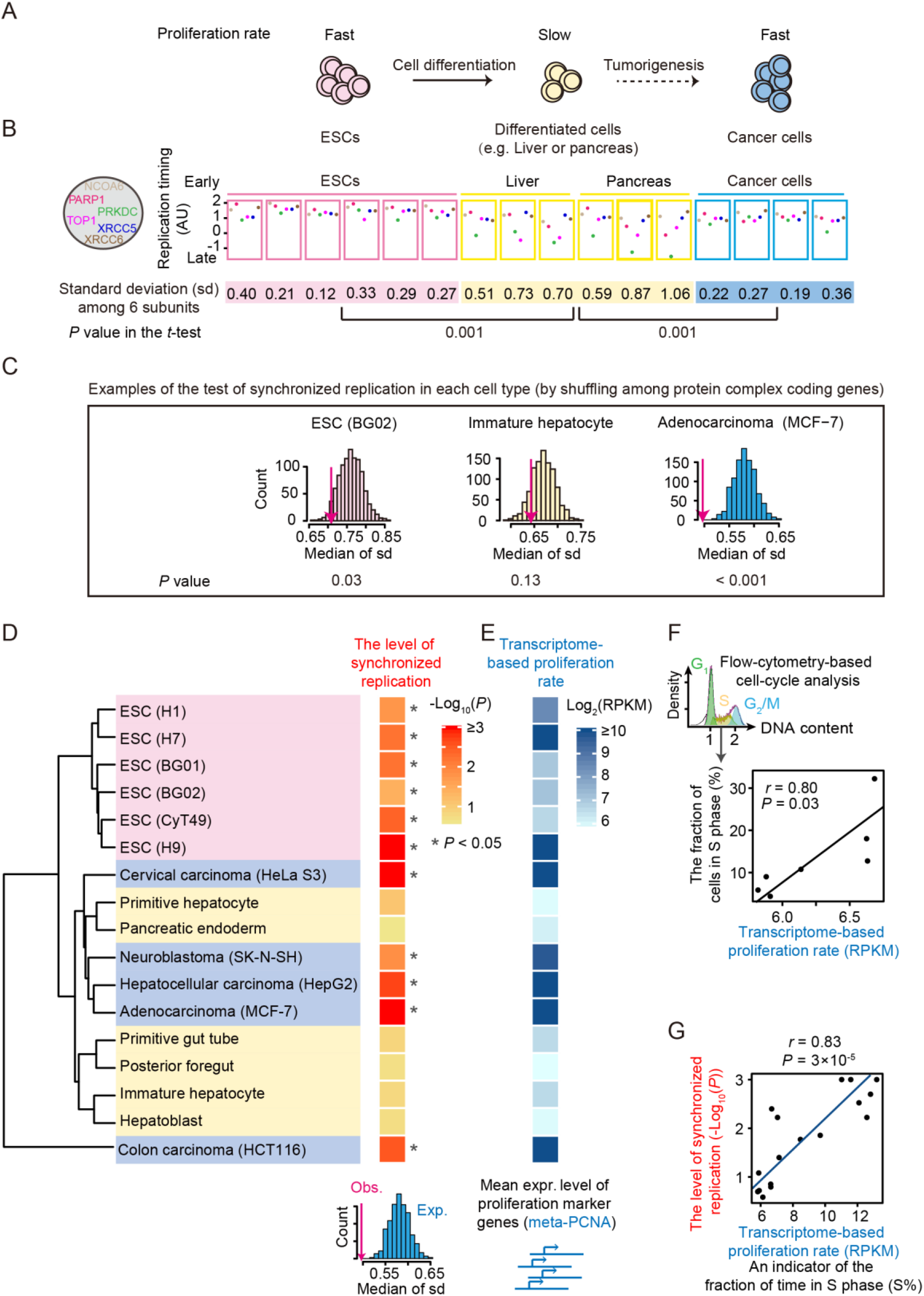
Genes in the same protein complex are replicated simultaneously only in fast-proliferating cells. (A) A schematic diagram shows the changes in cell proliferation rate during cell differentiation and tumorigenesis. (B) Replication timings of 6 genes encoding the same protein complex. Significant changes in standard deviation were estimated from *P* values in the *t*-tests. (C) Three examples of the tests for synchronized replication. (D) Synchronized replication occurs exclusively in ESCs and cancer cells. The dendrogram (left) shows the clustering of 14 cell lines/types based on the replication-timing profile of all genes in the genome. The heat map on the right shows the level of synchronized replication estimated from the permutation test. The asterisk represents significant synchronized replication (*P* < 0.05) in the corresponding cell line/type. (E) The proliferation rate was estimated from the average expression level of 11 proliferation marker genes (meta-PCNA) in the mRNA-seq data. (F) The fraction of cells in S phase (estimated by flow-cytometry) can be predicted from the transcriptome-based proliferation rate. Therefore, the transcriptome-based proliferation rate can be used as an indicator of the fraction of time in S phase (S%). (G) The proliferation rate and the level of synchronized replication are positively correlated.

We hypothesized that the loss of synchronized replication in differentiated cells was caused by the reduced power of natural selection for synchronized replication in slow-proliferating cells that spend a greater fraction of time in G0 phase and a smaller fraction of time in S phase (S%, **Fig. 1B**). To test this hypothesis, we estimated the proliferation rate from the average expression level of 11 proliferation marker genes (such as proliferating cell nuclear antigen, *PCNA*) for each cell line/type (**Fig. 3E**) (29). As expected, the proliferation rate was positively correlated with the proportion of cells in S phase among the 7 cell lines/types where flow-cytometry data were available (*r* = 0.8, *P* = 0.03, Pearson’s correlation, **Fig. 3F**) (27). The proliferation rate was positively correlated with the level of synchronized replication among 17 cell lines/types (*r* = 0.83, *P* = 3×10^−5^, Pearson’s correlation, **Fig. 3G**), suggesting an S%-dependent optimization of the replication-timing program in human cells.

### Synchronized replication is evolved convergently during tumorigenesis

Since cancer cells are “evolved” from various differentiated cells rather than ESCs. Consistently, ESCs and cancer cells did not form a monophyletic group (**Fig 3D**) when we clustered cell lines/types based on the similarity of their replicating-timing programs. Instead, four out of the five cancer cell lines were clustered with primitive hepatocytes and pancreatic endoderm cells, echoing their evolutionary origin of differentiated cells. Nevertheless, the replication-timing programs of these 4 cancer cell lines permit synchronized replication whereas those of primitive hepatocytes and pancreatic endoderm cells do not (**Fig. 3D**). More intriguingly, a human colon cancer cell line, HCT116, has a very different replication-timing program from all other cell types but exhibits the pattern of synchronized replication (**Fig. 3D**). Collectively, these observations suggest convergent cellular evolution during tumorigenesis to optimize the replication-timing program for fast proliferation.

### Differentiated cells lose synchronized replication mainly through a replication delay

To investigate the molecular mechanism by which synchronized replication is lost in differentiated cells, we identified 165 protein complexes in which the standard deviation of replication timing among subunits was significantly increased during differentiation (*P* < 0.05 in the *t*-tests, an example is shown in **Fig. 3B**). For each of the 491 genes encoding these protein complexes, we calculated the average replication timing among ESCs and differentiated cells, respectively (**Fig. 4A**). Among these genes, 92% exhibited a delay in replication in differentiated cells (top in **Fig. 4B**), likely through a postponement of firing time of the closest replication origins (an example is shown in **Fig. 4C**). In contrast, only 73% of non-complex encoding genes exhibited delays in replication timing (top in **Fig. 4B**, odds ratio = 4.1, *P* < 2.2×10^−26^, the Fisher’s exact test), suggesting that the differentiated cells lose synchronized replication preferentially through a replication delay.

**Figure 4.**
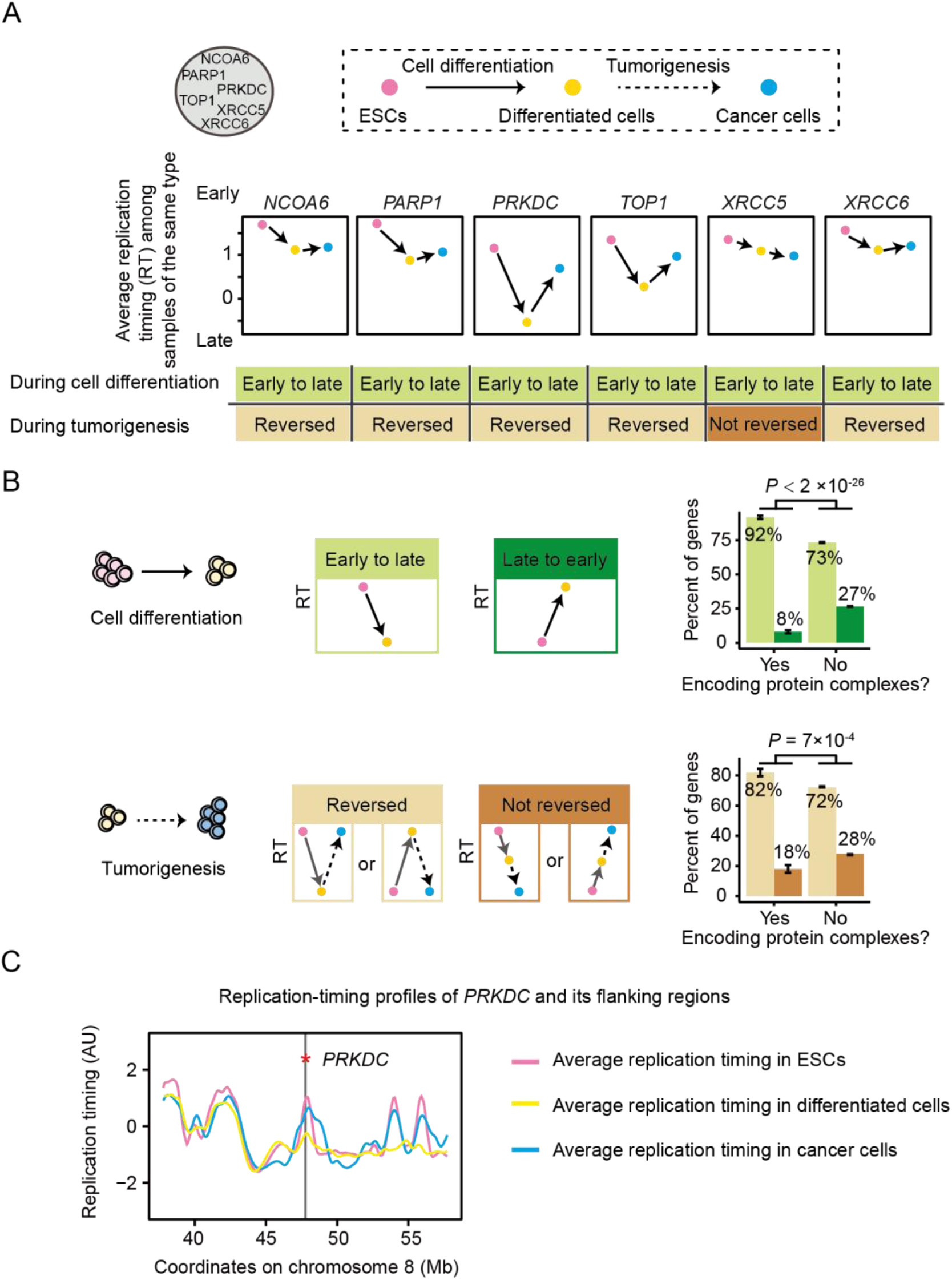
Mechanisms by which synchronized replication is lost and restored. (A) Calculation of the average replication timing (RT) among cells in the same group (ESCs, differentiated cells, or cancer cells). (B) The fraction of genes in each category during cell differentiation or tumorigenesis. (C) An example of changes in the replication-timing program. The loess-smoothed curves of replication timing in each cell group are shown. The vertical line and the asterisk indicate the position of the gene.

### Cancer cells reverse the change in replication timing during cell differentiation to restore synchronized replication

Among the 165 protein complexes in which synchronized replication is lost in differentiated cells, 79 protein complexes restored synchronized replication in cancer cells (bottom in **Fig. 4B**, *P* < 0.05 in the *t*-tests for the standard deviations between differentiated and cancer cells; an example is shown in **Fig. 3B**). Presumably, the restoration could occur either through i) reversing the change in replication timing during cell differentiation or ii) through an intergenic suppression that the replication timing of a second gene in the same protein complex follows that of the first. We found that during tumorigenesis, 82% genes reversed the changes during cell differentiation (an example is shown in **Fig. 4C**), significantly higher than the fraction (72%) among genes not encoding protein complexes (bottom in **Fig. 4B**, odds ratio = 1.7, *P* = 7×10^−4^, the Fisher’s exact test). Consistently, when we clustered the 17 cell types/lines with the replication timing of the genes encoding these 79 protein complexes, ESCs and cancer cells become closer in the dendrogram (**Fig. S3**).

## DISCUSSION

Abnormal replication-timing programs have been known to be related to disease and cancer (30–32). Our study provides a mechanism why a proper regulation of the replication-timing program is essential, especially for fast-proliferating cells: to maintain the dosage balance between early and late-replicating genes during S phase. We observed a convergent cancer evolution of replication-timing program toward ESCs, echoing previous analyses on the evolution of tumor cells at different levels, such as those at the transcriptome or the amino acid usage level (33–35).

We showed that the demand for dosage balance during S phase could cause synchronized replication of genes encoding the same protein complex. However, such synchronized replication could also have evolved under other selection pressures. For example, it may evolve to meet the demand for similar expression levels of genes encoding the same protein complex because replication timing is associated with gene expression level (36, 37). Nevertheless, this mechanism cannot explain why synchronized replication is lost in differentiated cells, where the genes encoding protein complexes remain expressed (**Fig. S4**) and the dosage balance among subunits remains important (**Fig. S5**). The selection for dosage balance during S phase uniquely predicts the loss of synchronized replication in slow-proliferating cells and the re-gain of it in cancer cells.

The synchronized replication of complex members is restored in cancer cells, although the number of mutations bared in each cancer cell is usually small (38). It is therefore unlikely that each complex restores the synchronized replication through individual mutations on its members during tumorigenesis. Master regulators of the replication-timing program exist which control the firing of multiple replication origins (39). In principle, the “switching” back of such master regulators to the ESC status could make cancer cells rapidly restored synchronized replication. Consistently, the re-gain of synchronized replication in cancer cells is mainly through reversing the changes in replication timing during cell differentiation (bottom in **Fig. 4B**).

Whereas the dosage imbalance during S phase can be partly relieved by the H3K56ac-associated transcription repression of newly replicated genes in yeast, the balance is not completely restored; early-replicating genes still exhibit a ~20% higher expression level during the mid-S phase (18). In cancer cells, the challenge of the dosage imbalance during S phase is likely greater, for two reasons. First, H3K56ac may play a less important role in repressing the expression of newly replicated genes in mammalian cells (21). Second, S phase lasts for a much longer time in mammalian cells than in yeast. For example, a HeLa cell divides every 24 hours and its S phase lasts for ~8 hours (40); HeLa cells need to suffer from the imbalance between early- and late-replicating genes for a few hours every 24 hours. By contrast, yeast has adapted to the life cycle of 24~48 hours per generation in the fermentation industry (41) or in nature which can be mimicked by a synthetic oak exudate medium (42, 43). However, the S phase lasts for less than 1 hour even in a poor carbon source (44), likely because the total time for DNA replication is mainly determined by the elongation rate of the DNA polymerase. The imbalance between early and late-replicating genes lasts for only dozen of minutes every 24~48 hours in yeast. Collectively, the natural selection for synchronized replication is likely stronger in cancer cells.

Some of the genes encoding the same protein complex form clusters on chromosomes (25, 26). This observation is often explained by the demand for coordinated gene expression (26), reduction in expression noise (45), and by the positive epistatic relationship among these genes (46). We showed that the synchronized replication hypothesis remained supported after controlling for such gene clusters (**Fig. S1**), yet the gene cluster itself could, in turn, be an evolutionary outcome of the selection for the dosage balance during S phase. Replication origins fire stochastically at the single-cell level (47). Therefore, the synchronization of replication timing is not robust in individual cells when these genes are interspersed in the genome and use the different replication origins. Forming a gene cluster is a more robust strategy for maintaining a dosage-sensitive relationship among genes.

Our results also have implications for DNA sequence evolution. For example, since replication timing is a major determinant of mutation rate (48, 49), we predict that genes encoding the same protein complex will have similar mutation rates due to synchronized replication. Consistently, it was reported that the evolutionary rate coevolves between a pair of genes that share a biological function or are co-expressed (50). Collectively, our study not only identifies the driving forces underlying the evolution of the replication-timing program but also provides new insights into the evolution of DNA sequences.

## METHODS

### Data retrievals

The information of 1,521 protein complexes in humans was downloaded from the Human Protein Reference Database (HPRD) release 9 (www.hprd.org). Among them, 1,317 were annotated completely and were used in this study. Chromosomal locations of these genes were retrieved from Ensembl release 87 (www.ensembl.org).

The replication-timing profiles used in this study were downloaded from the ReplicationDomain database (24) (https://www2.replicationdomain.com/) and are listed in **Table S1**. Below we briefly describe how the experiments were done to obtain the replication timing data. Detailed methods are in two previous studies (51, 52). Growing cells were pulse-labeled with bromodeoxyuridine (BrdU) for 1-2 hours, fixed, and then labeled with propidium iodide. Labeled cells were separated by fluorescence-activated cell sorter (FACS) into early and late-S bins. DNA with BrdU-incorporation was immunoprecipitated, differentially labeled, and co-hybridized to a microarray. The log_2_-transformed (early/late) ratio of the intensity of each probe was used to generate a replication-timing profile for the entire genome.

Gene expression data used in this study were downloaded from NCBI Gene Expression Omnibus (GEO) and are listed in **Table S1**.

### Estimation of gene replication timing

The list of 19,805 protein-coding genes was retrieved from human GRCh38.p12 annotation file that was downloaded from Ensembl (http://asia.ensembl.org/index.html). The average replication-timing ratio of probes having overlap with each gene was defined as the replication timing of this gene.

### Estimation of the proliferation rate with gene expression profiles

PCNA is a component of DNA polymerase δ and its expression level is a reporter of DNA synthesis. We defined a panel of 11 meta-PCNA genes whose expression is positively correlated with *PCNA*. Specifically, we normalized the average expression level of genes (log_2_(RPKM+1)) in 17 cell lines/types and calculated the Pearson’s correlation between PCNA and each of 131 previously identified candidate meta-PCNA genes. Genes with correlation coefficients greater than 0.9 were defined as meta-PCNA genes in this study (*PCNA, ZWINT, RFC3, LBR, TFDP1, SNRPB, SMC4, NUSAP1, BIRC5, UBE2C*, and *TROAP*). The proliferation rate was inferred from the average expression level (log_2_(RPKM+1)) of the 11 meta-PCNA genes (29).

### Estimation of the fraction of cells in S phase with flow-cytometry

Flow-cytometry data of propidium iodide-stained cells in 7 cell lines/types (listed in **Table S1**) were generated in a previous study (27). We estimated the fraction of cells belongs to one of the three stages in the cell cycle (G_1_, S, and G_2_/M) using FlowJo.

### Clustering of cell lines/types

The clustering of cell lines was performed with the function hclust in *R*. The Ward’s method was used.

### Loess smoothed replication-timing profiles

We combined the log_2_-transformed ratios (early/late) of the intensity of each probe for each of the three cell types (ESCs, differentiated cells, and cancer cells). The function loess.smooth in *R* was used to generate a smoothed profile (**Fig. 4C**) with the parameters span = 1/200 and evaluation = 2,000.

### Code availability

All codes to analyze the data and generate figures are available at https://github.com/YingChen10/Synchronized-replication-during-S-phase.

## Supporting information

## AUTHOR CONTRIBUTIONS

Y. C., K. L. and W. Q. conceived the research; Y. C., K. L., and X. C. analyzed the data; Y. C., L. B. C., and W. Q. wrote the manuscript.

## ACKNOWLEDGMENTS

We thank Tokyo Sasaki and David M Gilbert for providing the flow-cytometry data of cell-cycle analyses. We thank Zeyu Zhang for helpful discussion and Xionglei He and Mengyi Sun for critical comments on the manuscript. This work was supported by grants from the National Natural Science Foundation of China to W.Q. (91731302).

