## Supplementary material for "Selection for synchronized replication of genes encoding the same protein complex during tumorigenesis"

### SUPPLEMENTAL TABLE

**Table S1. Data sources of 17 cell lines/types used in this study[1].**

| Cell line | Replication timing | Gene expression | FACS for S% |
|---|---|---|---|
| ESC (BG01) | Int61646100 | GSE72923 | |
| ESC (BG02) | Int87960943 | GSE72923 | |
| ESC (CyT49) | Int83608519 | GSE63592 | Yes |
| ESC (H1) | Int79529114 | GSE67915 | |
| ESC (H7) | Int97847336 | GSE52657 | |
| ESC (H9) | Ext29405702 | GSE67915 | |
| Immature hepatocyte | Int54607190 | GSE63592 | Yes |
| Hepatoblast | Int79325459 | GSE63592 | Yes |
| Primitive hepatocyte | GSE63428 | GSE63592 | Yes |
| Primitive gut tube | GSE63428 | GSE63592 | Yes |
| Posterior foregut | GSE63428 | GSE63592 | Yes |
| Pancreatic endoderm | Int56654336 | GSE63592 | Yes |
| Adenocarcinoma (MCF−7) | Int67688290 | GSE33480 | |
| Colon carcinoma (HCT116) | Int90617792 | GSE33480 | |
| Cervical carcinoma (HeLaS3) | Int95117837 | GSE33480 | |
| Neuroblastoma (SK-N-SH) | Int67184500 | GSE90265 | |
| Hepatocellula carcinoma (HepG2) | Int49277082 | GSE87999 | |

[1] Int* and Ext* are accession numbers in the Replication Domain database (https://www2.replicationdomain.com/). GSE* are accession numbers in the NCBI Gene Expression Omnibus (GEO; http://www.ncbi.nlm.nih.gov/geo/).

### SUPPLEMENTAL FIGURES

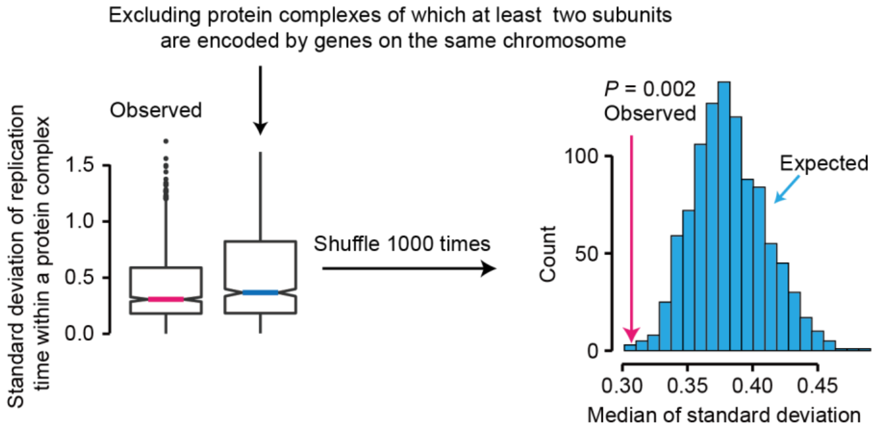


**Figure S1.** The observed standard deviation of replication timing within a protein complex is significantly smaller than the random expectation after we excluded protein complexes of which at least two subunits are encoded by genes on the same chromosome.

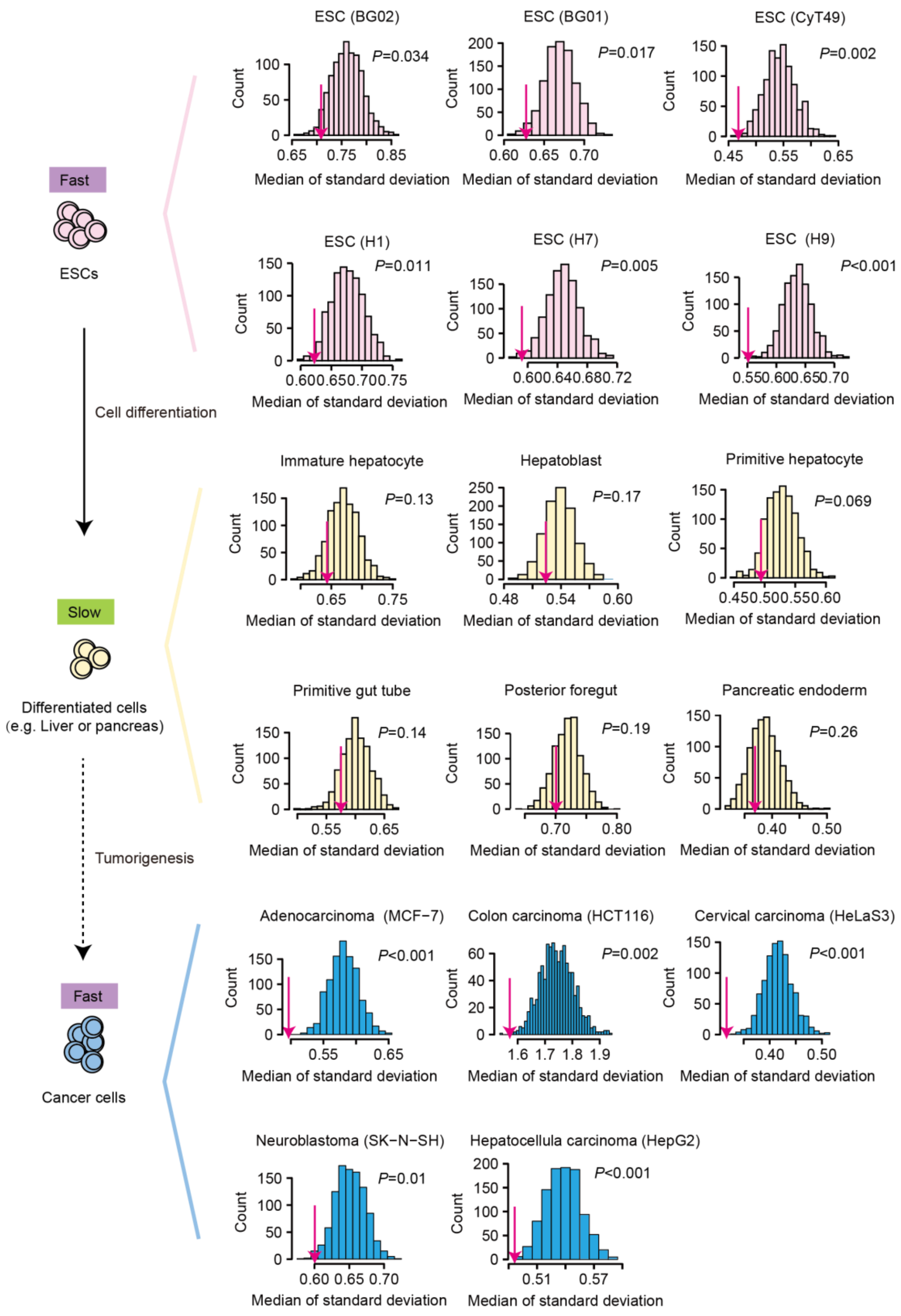


**Figure S2. Determination if synchronized replication occurs in each of the 14 cell types.** The observed median of the standard deviation of replication timing within a protein complex for each cell type (magenta arrows) is shown on the distribution of the random expectation of 1,000 shuffling. Genes encoding protein complexes were shuffled (as in Fig. 2C).

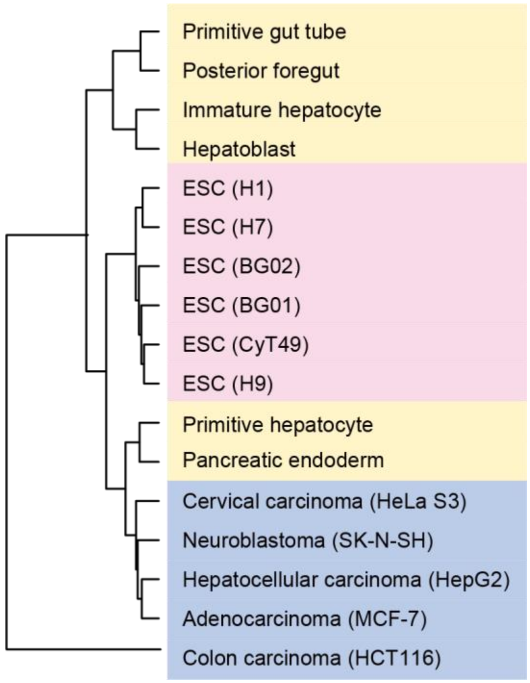


**Figure S3.** The dendrogram shows the clustering of 17 cell lines/types based on the replication-timing profile of genes encoding 79 protein complexes which restore dosage balance in cancer cells. Similar to Fig. 3D.

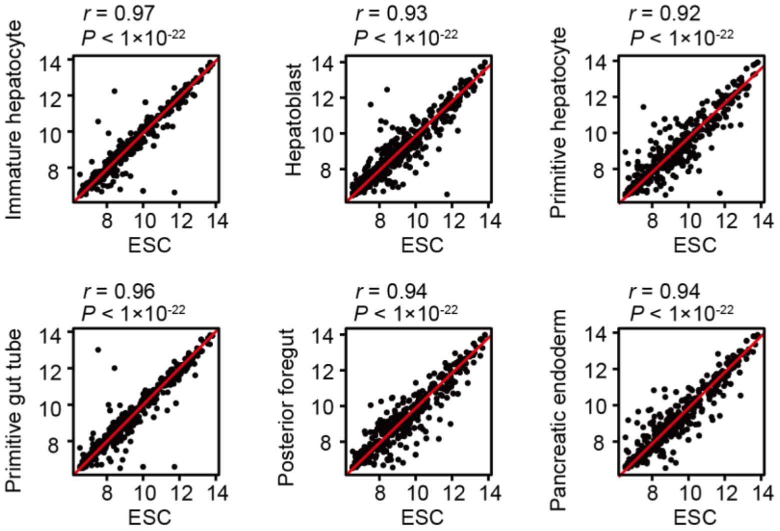


**Figure S4. Expression levels are highly correlated between ESCs and each of the 6 differentiated cells.** Each dot represents one of the 491 genes encoding the 165 protein complexes in which the standard deviation of replication timing was significantly increased in differentiated cells. Pearson's correlation coefficients, *r*, and the corresponding *P* values are shown.

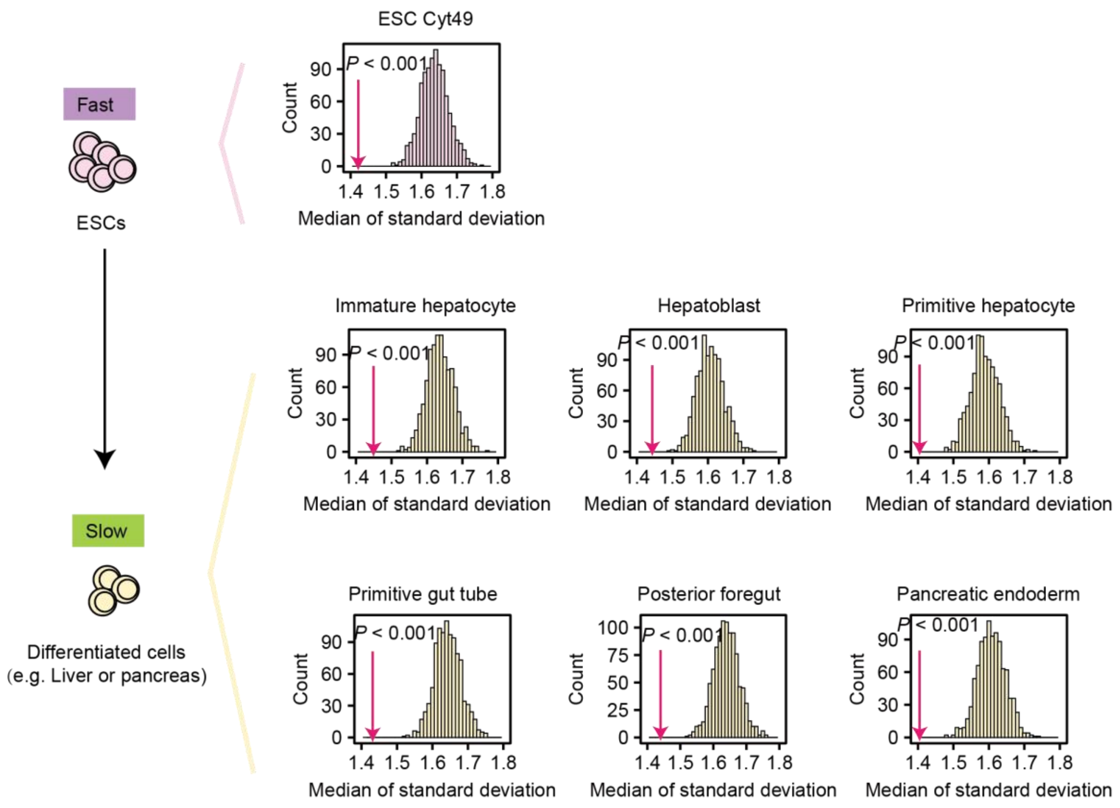


**Figure S5. Dosage balance among members of the same protein complex remains important in 6 differentiated cell types.** For each protein complex, we calculate the standard deviation of the expression level ($\log_2$(RPKM+1)) of genes encoding the protein complex (the pink arrows). We shuffled the expression level of the protein complex-coding genes 1000 times to generate a distribution of random expectation. The median of the observed standard deviation was significantly smaller than the random expectation in all 7 cell types, indicating the dosage balance remains importance in differentiated cells. Note that we assumed that genes encoding the same protein complex have a stoichiometric relationship of 1:1 because the stoichiometric data of protein complexes are not completely available. Nevertheless, our test here was conservative because the observed standard deviation would be overestimated if the actual stoichiometric relationship was not 1:1.
